# The influence of early moderate prenatal alcohol exposure and maternal diet on offspring DNA methylation: a cross-species study

**DOI:** 10.1101/2023.06.15.544516

**Authors:** Mitchell Bestry, Alexander N. Larcombe, Nina Kresoje, Emily K Chivers, Chloe Bakker, James P Fitzpatrick, Elizabeth J Elliott, Jeffrey M Craig, Evelyne Muggli, Jane Halliday, Delyse Hutchinson, Sam Buckberry, Ryan Lister, Martyn Symons, David Martino

## Abstract

Alcohol consumption in pregnancy can affect genome regulation in the developing offspring but results have been contradictory. We employed a physiologically relevant murine model of short-term moderate prenatal alcohol exposure (PAE) resembling common patterns of alcohol consumption in pregnancy in humans. Early moderate PAE was sufficient to affect site-specific DNA methylation in new-born pups without altering behavioural outcomes in adult littermates. Whole-genome bisulfite sequencing of neonatal brain and liver revealed stochastic influence on DNA methylation that was mostly tissue-specific, with some perturbations likely originating as early as gastrulation. DNA methylation differences were enriched in non-coding genomic regions with regulatory potential indicative of broad effects of alcohol on genome regulation. Replication studies in human cohorts with fetal alcohol spectrum disorder suggested some effects were metastable at genes linked to disease-relevant traits including facial morphology, intelligence, educational attainment, autism, and schizophrenia. In our murine model, a maternal diet high in folate and choline protected against some of the damaging effects of early moderate PAE on DNA methylation. Our studies demonstrate that early moderate exposure is sufficient to affect fetal genome regulation even in the absence of overt phenotypic changes and highlight a role for preventative maternal dietary interventions.

## Introduction

Alcohol consumption in pregnancy is a leading cause of neurodevelopmental impairments in children (1). Alcohol can pass through the placenta acting as a teratogen in fetal tissues causing physical, cognitive, behavioural, and neurodevelopmental impairment in children at high doses with lifelong consequences for health (2). Fetal Alcohol Spectrum Disorder (FASD) and Fetal Alcohol Syndrome (FAS) can arise at binge levels of exposure, although not always at lower levels of exposure. Whether PAE is sufficient to induce overt physiological abnormalities depends on multiple environmental and genetic factors including the dose and timing of alcohol use during pregnancy, maternal diet, smoking, stress, and potentially other factors that collectively influence fetal outcomes (2–4).

Patterns of alcohol consumption in pregnancy vary, but epidemiological surveys suggest most women in Western countries consume low to moderate levels between conception until recognition of pregnancy (5), after which time consumption largely ceases, apart from occasional use (6). While the effects of binge levels of exposure are well documented as able to cause FASD, more subtle effects that reflect the more common patterns of drinking are unclear and more research is needed to support public health initiatives to reduce alcohol consumption in pregnancy.

Studies suggest alcohol can disrupt fetal gene regulation through epigenetic mechanisms (7). DNA methylation is one epigenetic mechanism involving the catalytic addition of methyl groups to cytosines bases within cytosine-guanine (CpG) dinucleotide motifs during one-carbon metabolism. Methylation of DNA can alter chromatin density and influence patterns of gene expression in a tissue-specific and developmentally appropriate manner and disruption to this process may cause some of the difficulties experienced by people with FASD (8, 9). Previous studies on human participants (10–13) and animals (14–16) report that alcohol can disrupt DNA methylation either globally (12–15), and/or at specific gene regions (11, 15, 16). Our recent systematic review, however, found limited replication of effects between studies suggesting the effects of alcohol on DNA methylation may be stochastic and influenced by numerous confounding factors (7). PAE can either directly inhibit DNA methyltransferase enzymes or disrupt one-carbon metabolism via inhibition of bioavailability of dietary methyl donors, such as folate and choline to the fetus (17, 18). Choline, in particular, has been explored in several clinical trials to reduce cognitive deficits caused by PAE in affected individuals (19, 20), or when administered during pregnancy (21, 22), with results suggesting a high methyl donor diet (HMD) could at least partially mitigate the adverse effects of PAE on various behavioural outcomes.

Given the lack of clarity around the effects of typical patterns of alcohol consumption, which often do not cause observable phenotypes, we conducted an epigenome-wide association study of early moderate PAE in mice. Regions identified as sensitive to gestational alcohol exposure were replicated in human cohorts. The study was a controlled intervention investigating the impact of early moderate PAE on offspring DNA methylation comparing exposed and unexposed mice, with an additional arm comparing the effect of alcohol exposure in the context of a high methyl donor maternal diet. The exposure period covers the equivalent of pre-conception up until the first trimester in humans when neurulation occurs, reflecting a typical situation in which women may consume alcohol up until pregnancy recognition (5)(23). The primary outcome of the study was differences in offspring DNA methylation and secondary outcomes of behavioural deficits across neurodevelopmental domains relevant to FASD were also examined. We employed whole-genome bisulfite sequencing (WGBS) for unbiased assessment of CpG DNA methylation in newborn brain and liver, two target organs affected by ethanol (24), to explore tissue-specificity of effects and to determine any ‘tissue agnostic’ effects which may have arisen prior to the germ-layers separating in early gastrulation. We also conducted candidate gene testing of regions identified in prior studies as sensitive to early moderate PAE. Our study provides cogent evidence that common patterns of drinking can have measurable effects on fetal gene regulation, highlighting a role for maternal dietary support in public health interventions.

## Results

### Comparison of prenatal characteristics across treatment groups

To investigate the effects of early moderate PAE and a high-methyl donor diet (HMD) across pregnancy on offspring DNA methylation and behavioural outcomes, we employed a murine model with four treatment groups. Treatment groups were designed to assess the effect of alcohol on offspring DNA methylation compared to control mice. An additional treatment group included a maternal diet high in methyl donors, with and without alcohol exposure (Figure 1).

**Figure 1.**
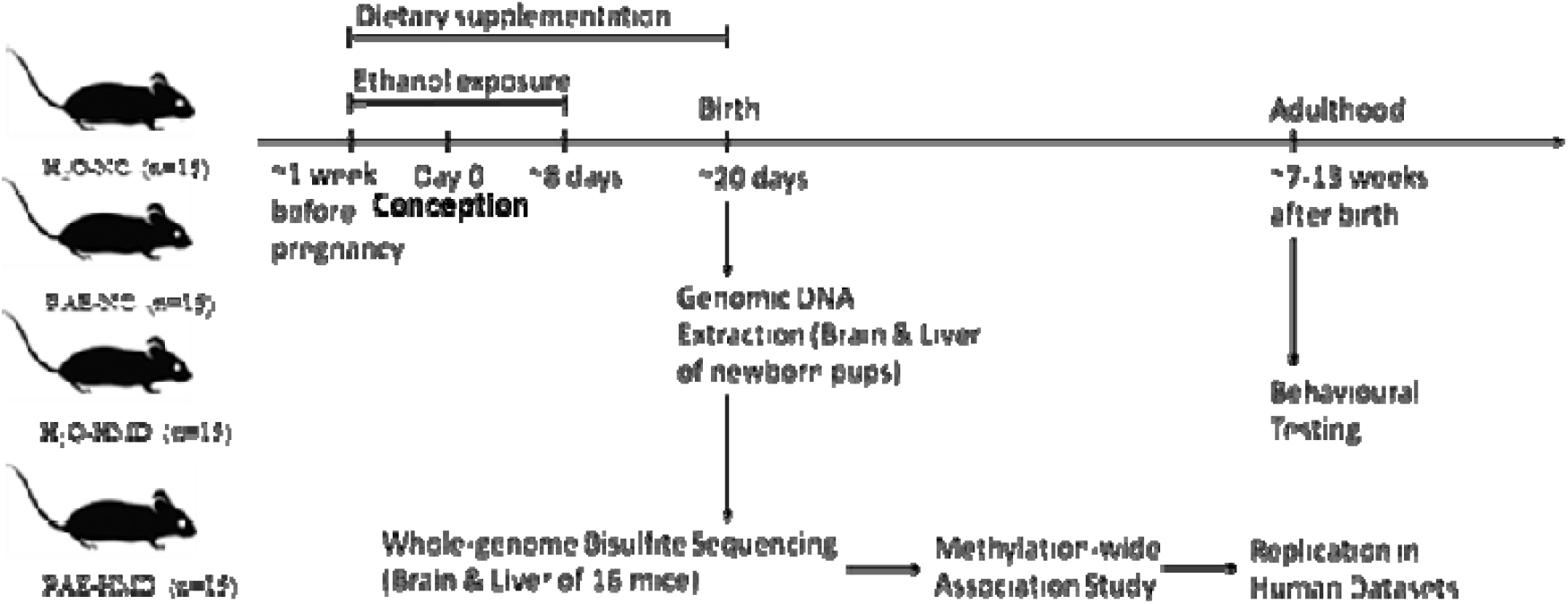
Overview of PAE model. A schematic representation of the experiment design is shown in figure 1. Fifteen dams were allocated to each treatment group. Prenatal alcohol exposure (PAE) mice were exposed to ethanol (10% v/v in non-acidified reverse osmosis drinking water *ad libitum*) from one week before pregnancy to gestational days 8-10 and the remaining mice received water (H_2_0). The PAE and H_2_0 groups received either normal chow (NC) or a high methyl donor (HMD) diet (NC containing 20 mg/kg folate and 4970 mg/kg choline) from one week before pregnancy until birth.

The trajectory of weight gain during pregnancy was similar across all treatment groups with some evidence of more rapid weight gain in the HMD groups in the last 2 days (Linear mixed effects regression model; H_2_0-HMD: -2.282 ± 0.918 g, P = 0.0177; PAE-HMD: -1.656 ± 0.814 g, P = 0.0493; figure S1a), although the total amount of weight gained between days 1 and 17-19 was not significantly different between treatment groups by linear mixed effects regression (Figures S1a-b). The total amount of liquid consumed over the course of pregnancy was significantly lower in HMD dams by unpaired t-test (Figures S1b and S2c). There was no significant difference in the average litter size (6.525 ± 0.297 pups, figure S1c) and pup sex ratios (Figure S1d) between treatment groups by unpaired t-test.

### Effects of early moderate PAE on DNA methylation in newborn mice tissue

A subset of 16 newborn pups (n = 4 per treatment) matched for sex and litter size were selected for whole-genome bisulfite sequencing. Both neonatal brain and liver were harvested to investigate the effects of early moderate PAE on fetal CpG DNA methylation. A total of 21,842,961 CpG sites were initially available for analysis.

Global levels of DNA methylation stratified across different genomic contexts were preserved across treatment conditions, with no major differences in average DNA methylation content between groups (Figure S4). To investigate region-specific effects of early moderate PAE on newborn DNA methylation, we conducted genome-wide testing comparing exposed and un-exposed mice on the normal chow. We identified 78 differentially methylated regions (DMRs) in the brain and 759 DMRs in the liver (P < 0.05 and mean difference in methylation across the DMR with PAE (delta) > 0.05) from ∼19,000,000 CpG sites tested after coverage filtering (Figure 2a-b). These regions were annotated to nearby genes using *annotatr* and are provided in Tables S1-2. Two of the DMRs overlapped in mouse brain and liver (tissue agnostic), but the remainder were tissue specific. Among these tissue agnostic regions was the *Impact* gene on chromosome 18, which had lower methylation in PAE+NC mice compared to H_2_0+NC mice in both the brain and liver (Figure 2c-d). The other tissue agnostic region was within 5kb downstream of *Bmf* and had higher DNA methylation in brain and liver tissue of PAE+NC mice.

**Figure 2.**
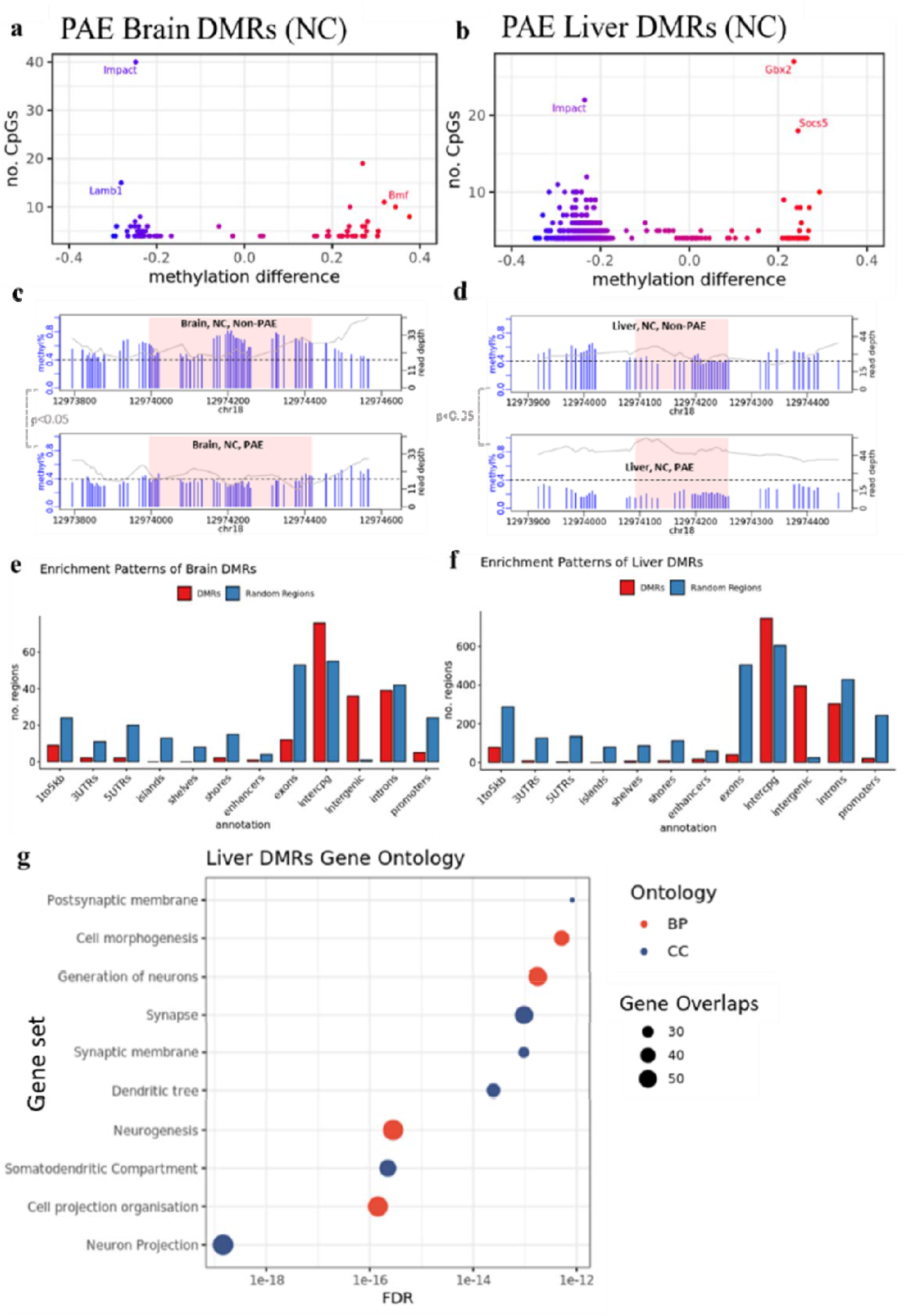
PAE was associated with site-specific differences in offspring DNA methylation. The majority of DMRs lost methylation with PAE in (a) brain and (b) liver of mice given normal chow. Each point represents one DMR. Point colour indicates change in DNA methylation with PAE. PAE was also associated with lower methylation in the DMRs identified in the promoter of the *Impact* gene in (c) brain and (d) liver, within NC mice. Each plot represents a separate treatment group. Each blue vertical line indicates a CpG site, with the height and corresponding left y-axis indicating the methylation ratio. The grey line and corresponding right y-axis indicate coverage at each CpG site. The black horizontal dotted line indicates 40% methylation for comparison purposes. The x-axis indicates the base position on chromosome 18, with the pink shaded area highlighting the DMR. DMR plots include 200 base pair flanking regions on each side of the DMR. DMRs identified in (e) brain and (f) liver were enriched in intergenic and inter-CpG regions, whilst being underrepresented in CpG and gene regions. The bar plot compares the number of WGBS DMRs in red to a set of equivalent randomly generated regions in blue. (g) Gene ontology analysis of liver DMRs shows enrichment within neuronal cellular components and biological processes. BP/red point = biological process, CC/blue point = Cellular component. X-axis of point indicates FDR of ontology. Size of point indicates number of overlapping genes with ontology. There were insufficient number of DMRs identified in the brain for a gene ontology analysis.

Lower DNA methylation with early moderate PAE in NC mice was more frequently observed in liver DMRs (93.5% of liver DMRs), while brain DMRs were almost equally divided between lower and higher DNA methylation with early moderate PAE (52.6% of brain DMRs had lower DNA methylation with early moderate PAE). Some DMRs localised to the same genes in both brain and liver, although they were different regions. The three genes affected by PAE in both brain and the liver tissues were the Autism Susceptibility Gene 2 (*Auts2*), Androglobin (*Adgb*), and RNA Binding Protein Fox 1 (*Rbfox1*) genes (Table 1). In both brain and liver tissues, DMRs were enriched in non-coding intergenic and open sea regions and relatively underrepresented in coding and CpG island regions (Figure 2e-f). Using open chromatin assay and histone modification datasets from the ENCODE project, we found enrichment (p < 0.05) of DMRs in open chromatin regions (ATAC-seq), enhancer regions (H3K4me1), and active gene promoter regions (H3K27ac), in mouse fetal forebrain tissue and fetal liver (Table 2). Gene ontology enrichment analysis of liver DMRs that did localise to genes showed enrichment in ten predominantly neuronal pathways, with neuron projection being the most significant (Figure 2g, Tables S3-4).

**Table 1:** Table of DMRs identified in the intronic regions of genes that contained DMRs in both the brain and liver. Δmeth indicates the percentage change in average methylation level within the DMR with PAE compared to non-PAE mice.

| Gene | Tissue | Intronic DMR | Width | No. CpGs | Δmeth | P-value |
| --- | --- | --- | --- | --- | --- | --- |
| *Auts2* | Brain | chr5:131510296-131510465 | 170 | 5 | -23.5% | <0.05 |
| *Auts2* | Liver | chr5:131621828-131621999 | 172 | 4 | -22.5% | <0.05 |
| *Adgb* | Brain | chr10:10455557-10455883 | 327 | 4 | -25.0% | <0.05 |
| *Adgb* | Liver | chr10:10353338-10353613 | 276 | 4 | -25.9% | <0.05 |
| *Rbfox1* | Brain | chr16:6813039-6813217 | 179 | 5 | -24.3% | <0.05 |
| *Rbfox1* | Liver | chr16:6781985-6782330 | 346 | 5 | -22.6% | <0.05 |

**Table 2.** Number and percentage of brain and liver DMRs that overlap with tissue-specific regulatory regions. ATAC-seq, H3K4me1 and H3K27ac regions were obtained at 0 days postnatal from the ENCODE database. P-values for permutation testing using a randomisation strategy.

| Assay type | Brain DMRs | Brain randomised | Liver DMRs | Liver randomised |
| --- | --- | --- | --- | --- |

|  |  | regions |  | regions |
| --- | --- | --- | --- | --- |
| ATAC-seq | 21/78 (26.92%),<br>P = 0.01 | 1/78 (1.28%), P =<br>0.16 | 53/759 (6.98%)<br>P = 0.01 | 22/759 (2.90%)<br>P = 0.31 |
| H3K4me1 | 4/78 (5.13%)<br>P = 0.03 | 2/78 (2.56%)<br>P = 0.18 | 38/759 (5.01%)<br>P = 0.05 | 35/759 (4.61%)<br>P = 0.32 |
| H3K27ac | 9/78 (11.54%)<br>P = 0.01 | 2/78 (2.56%)<br>P = 0.74 | 48/759 (6.32%)<br>P = 0.01 | 19/759 (2.50%)<br>P = 0.26 |

### HMD mitigates the effects of early moderate PAE on DNA methylation

To determine whether administration of a HMD throughout pregnancy could mitigate the effects of PAE on offspring DNA methylation, we examined alcohol sensitive DMRs identified in the previous analysis in the HMD mice. Compared to control mice (H20+NC), PAE+HMD mice exhibited significant (p < 0.05) DNA methylation differences in only 12/78 (15%) brain (Table S7), and 124/759 (16%) liver (Table S8) DMRs, suggesting the effects were predominantly mitigated. Effect sizes compared to mice on the normal chow were substantially lower, in some cases more than 25% reduced in mice on the high methyl donor diet (Figure 3).

**Figure 3.**
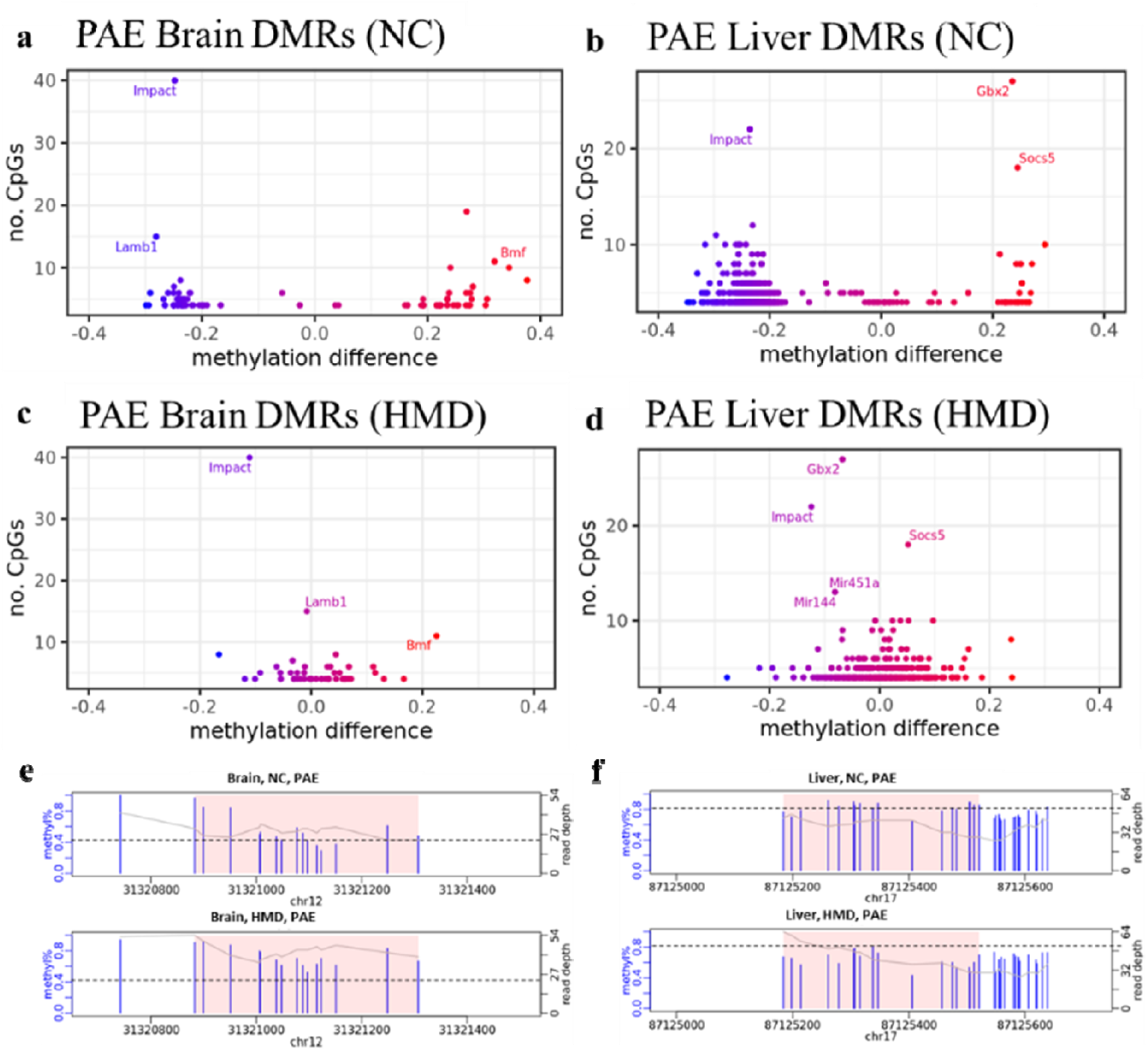
HMD partially mitigated effects of PAE on offspring DNA methylation. Average DNA methylation effect sizes above 30% with PAE were observed in some (a) brain, and (b) liver DMRs in NC mice. Mean absolute difference in methylation with PAE is reduced within the HMD mice in (c) brain, and (d) liver. Each point represents one DMR. Point colour indicates change in DNA methylation with PAE. Points with a high number of CpGs and methylation difference are annotated with associated gene if located within a genic region. HMD was associated with (e) higher methylation in the DMR identified proximal to *Lamb1* on chromosome 12 in brain and (f) lower methylation in the DMR identified proximal to *Socs5* on chromosome 17 in liver. Each plot represents a separate treatment group. Each blue vertical line indicates a CpG site, with the height and corresponding left y-axis indicating the methylation ratio. The grey line and corresponding right y-axis indicate coverage at each CpG site. The black horizontal line indicates (e) 40% and (f) 80% methylation for comparison purposes. The x-axis indicates the base position on the chromosome, with the pink shaded area highlighting the DMR. DMR plots include 200 base pair flanking regions on each side of the DMR.

### Effects of early moderate PAE and HMD on behavioural outcomes in adult mice

Remaining littermates from each treatment group were reared to adulthood and underwent behavioural testing assessing various neurocognitive domains that can be affected in FASD including locomotor activity, anxiety, spatial recognition, memory, motor coordination and balance. There was no evidence that early moderate PAE had a significant effect on any of the behavioural outcomes tested (Figure S3). Mice exposed to HMD exhibited greater locomotor activity, in terms of distance travelled (Figure 4).

**Figure 4.**
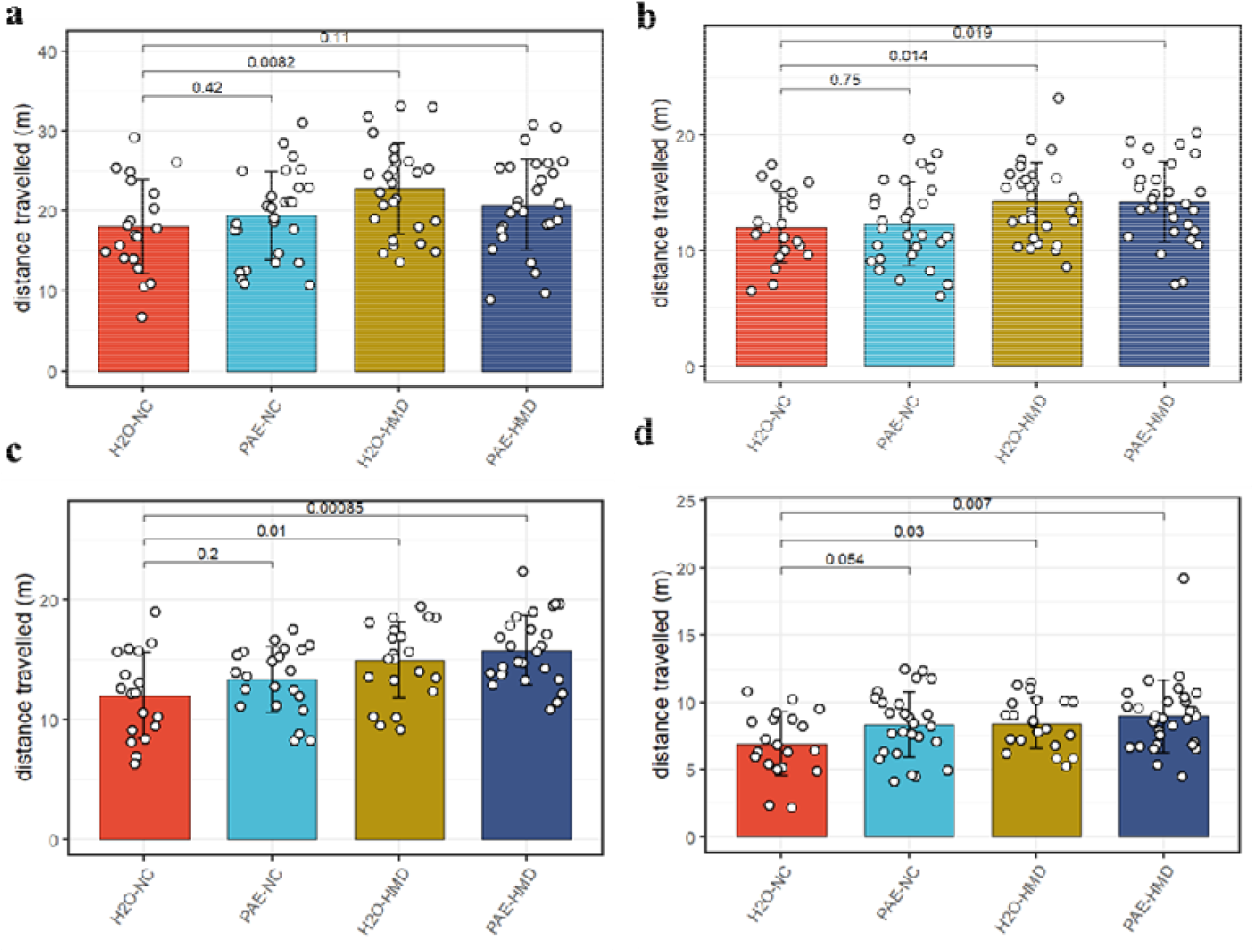
HMD was associated with increased locomotor activity. HMD was associated with increased locomotor activity compared to NC, indicated by significantly greater total distance travelled in the (a) open field test (N = 104), (b) object recognition test (N = 108), (c) elevated plus maze test (N = 88), and (d) object in place test (N = 98) by unpaired t-test. Bars show mean and standard deviation. Each point represents one mouse. NC = normal chow, HMD = high methyl diet, PAE = prenatal alcohol exposure. Time interval for each mouse was (a-c) 300 seconds and (d) 180 seconds.

### Replication studies in Human PAE and FASD case-control cohorts

We undertook validation studies by examining PAE sensitive regions identified in our murine model using existing DNA methylation data from human cohorts to address the generalizability of our findings. Only 36 of the 78 (46.2%) brain DMRs, and 294 of the 759 (38.8%) liver DMRs, had homologous regions in the human genome that were able to be tested. In this validation study, DNA methylation array data from 147 newborn buccal swabs from the Asking Questions About Alcohol in Pregnancy (AQUA) cohort (5) were available from this cohort (96 moderate PAE and 51 controls). We performed differential testing on a total of 1,898 CpG sites that corresponded to mouse DMRs, comparing ‘never exposed’ newborns to ‘any exposure’ and found no evidence of differential DNA methylation at these CpG (data not shown). We also accessed publicly available DNA methylation array measurements from buccal swabs taken from a Canadian clinical cohort of children with diagnosed FASD, and controls. To avoid confounding due to ancestry, we analysed the 118 Caucasian individuals (30 FASD and 88 controls). Differential testing of a total of 2,316 CpG sites that were homologous to mouse DMRs statistically replicated 7 DMR associations with FASD status (FDR P < 0.05) after adjusting for participant age, sex, array number, and estimated cell counts (Table 3). Visual comparisons of DNA methylation across these seven DMRs revealed striking differences in effect sizes between people with FASD and our murine model (Figure 5). Genes associated with these DMRs are linked to clinically relevant traits in the GWAS catalogue including facial morphology (*GADD45A* (26)), educational attainment (*AP2B1* (27), intelligence (*RP9* (28)), autism and schizophrenia (*ZNF823* (29)).

**Figure 5.**
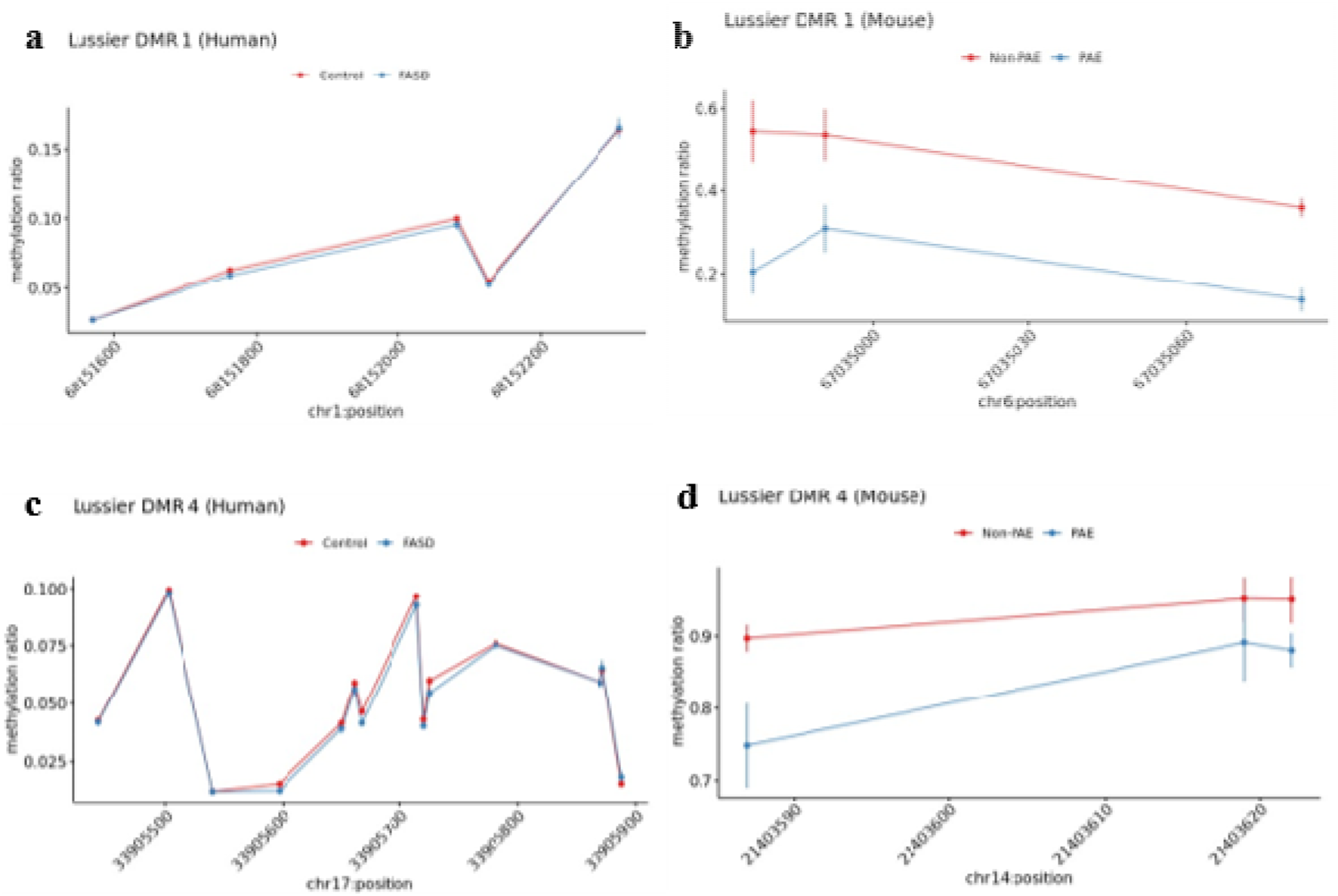
Seven PAE DMRs identified in the murine model were successfully replicated in the Lussier et al. human FASD cohort. Examples of two PAE DMRs that had significantly lower DNA methylation with a clinical diagnosis of FASD in the Lussier et al. cohort (a and c), while their mouse liftover DMR also had significantly lower DNA methylation with PAE in the murine model experiment (b and d).

**Table 3.**
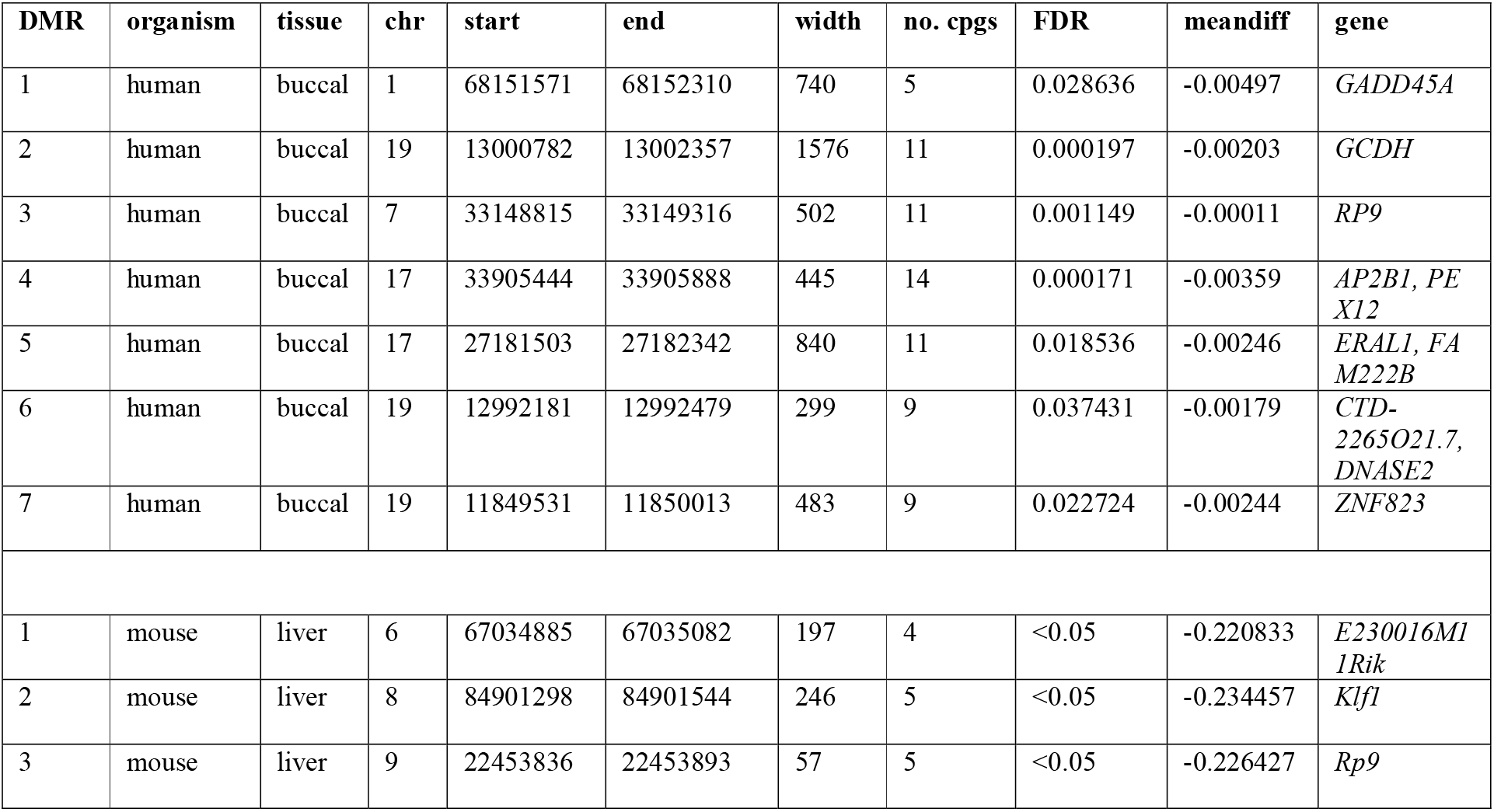
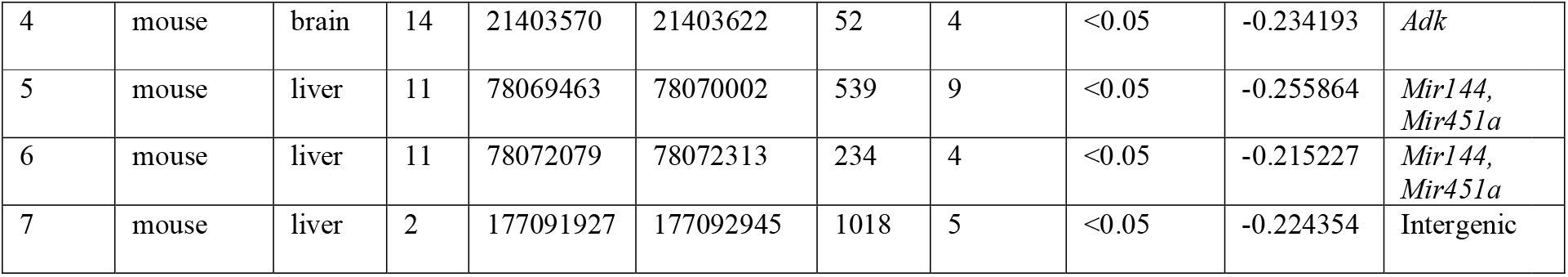
DMRs identified in the murine model that were validated in the Lussier et al. human case-control cohort for a clinical diagnosis of FASD. The upper section describes properties of Lussier et al. human DMRs. The lower section describes properties of the equivalent murine model DMRs.

### Candidate Gene Analysis of previously defined alcohol sensitive regions

We also undertook a replication analysis in our murine data of previously published alcohol sensitive regions by undertaking a systematic review of previously published mammalian studies (7). Candidate gene studies identified 21 CpG sites (FDR<0.05) in the brain from 15,132 CpG sites tested, including two sites in the *Mest* (*Peg1*) gene and 19 sites in *Kcnq1* (*KvDMR1*) (Table S9). There were nine FDR-significant CpG sites identified in the liver out of 15,382 CpG sites tested, all of which were in *Peg3* (Table S10). All FDR-significant CpG sites from both tissues had higher DNA methylation in mice with PAE.

## Discussion

In this study, we found that moderate early (first trimester) PAE was sufficient to induce site-specific differences to DNA methylation in newborn pups without causing overt behavioural outcomes in adult mice. Although global levels of DNA methylation were not significantly different with PAE, regional analysis demonstrated widespread effects characterised predominantly by lower DNA methylation with PAE, mostly at non-coding regions of the genome. In our model, alcohol effects on DNA methylation were predominantly tissue-specific, with only two genomic regions and four genes that were similarly affected in both liver and brain. These perturbations may have been established stochastically because of PAE to the early embryo and maintained in the differentiating tissue. Further analysis in different germ layer tissues is required to formally establish this. Indeed, most of the observed effects were tissue-specific, with more perturbations to the epigenome observable in liver tissue, which may reflect the liver’s specific role in metabolic detoxification of alcohol. Alternatively, cell type composition differences between brain and liver might explain differential sensitivity to alcohols effects. Generally, DMRs were enriched in non-coding regions of the genome with regulatory potential, suggesting alcohol has broad effects on genome regulation.

Both the human replication studies and the candidate gene analysis provide validity to our model for recapitulating some of the genomic disturbances reported in patients with clinical FASD. It is remarkable that some associations identified in our murine model of early moderate exposure were recapitulated in human subjects with FASD despite species and biosample differences, suggesting that at least some DNA methylation changes are stable over time. Notably the effect size for replicated regions were strikingly smaller in blood samples from subjects with FASD suggesting the dose and duration of exposure may need to exceed a high threshold to survive reprogramming in the blood. We speculate this may explain lack of reproducibility in the AQUA cohort.

In the candidate gene analysis we replicated previously published reports of decreased DNA methylation within *Peg3* and *KvDMR1* from South African children with fetal alcohol syndrome (11). Both genes are methylated in a parent-of-origin specific manner, suggesting that alcohol may affect imprinting processes. Previous rodent and human studies have identified DNA methylation differences with PAE in imprinted regions such as the *Igf2/H19* locus (13, 16, 30), although results are not entirely consistent (31, 32). On the balance of this, we speculate duration of exposure, dose, and other tissue-related factors all likely influence the extent to which genome-regulation is perturbed and manifests as differences in DNA methylation.

Our results are encouraging for biomarker studies and aid in the prioritisation of associations for future follow-up, particularly in relation to diagnosis of FASD. For example, three genes that were validated in the Lussier et al. cohort are zinc finger proteins (*RP9*, *PEX12*, and *ZNF823*) that play an important role in fetal gene regulation. Notably, *PEX12* is associated with Zellweger syndrome, which is a rare peroxisome biogenesis disorder (the most severe variant of Peroxisome biogenesis disorder spectrum), characterised by neuronal migration defects in the brain, dysmorphic craniofacial features, profound hypotonia, neonatal seizures, and liver dysfunction (33). Future studies could perform transcriptomic analysis to investigate Another key finding from this study was that HMD mitigated some of the effects of PAE on DNA methylation. Although a plausible alternative explanation is that some of the PAE regions were not reproduced in the set of mice given the folate diet, our data are consistent with preclinical studies of choline supplementation in rodent models (34, 35) (36). Moreover, a subset of PAE regions were statistically replicated in subjects with FASD, suggestive of robust associations. Although our findings should be interpreted with caution, they collectively support the notion that alcohol induced perturbation of epigenetic regulation may occur, at least in part, through disruption of the one-carbon metabolism. The most encouraging aspect of this relates to the potential utility for evidence-informed recommendations for dietary advice or supplementation, particularly in population groups with limited access to antenatal care or healthy food choices.

Strengths of this study include the use of controlled interventions coupled with comprehensive assessment of the effects of PAE on multiple tissues. We also performed whole-genome bisulfite sequencing representing the gold standard in DNA methylation analysis, which to our knowledge has not been performed before in the context of murine PAE studies. Our findings were partially generalisable in replication studies addressing the robustness of our experimental approach. Caveats of our study design include a limited ability to determine the contribution of specific cell types within tissues to the methylation differences observed, and we did not assess markers of brain or liver physiology. Additionally, we employed an ad-libitum alcohol exposure model rather than direct dosing of dams. Although the trajectories of alcohol consumption were not statistically different between groups, this introduces more variability into alcohol exposure patterns, and might impact offspring methylation data. Despite these limitations, the results were meaningful in the context of typical patterns of alcohol consumption in human populations.

In conclusion, this study demonstrates that early moderate PAE can disturb fetal genome regulation in mice and humans and supports current public health advice that alcohol consumption during pregnancy, even at low doses, may be harmful.

## Materials and Methods

### Murine subjects and housing

To study the effects of early moderate PAE on offspring DNA methylation processes, we adapted a murine model study design that has previously reported DNA methylation changes at the A^vy^ locus in Agouti mice (37) (Figure 1). This study received animal ethics approval from the Telethon Kids Institute Animal Ethics Committee (Approval Number: 344). Sixty nulliparous C57BL/6J female mice aged ∼8 weeks were mated with equivalent stud male mice. Pregnant dams were randomly assigned to one of four treatment groups (n = 15 dams per group) that varied based on composition of the drinking water and chow given to the dams:

i. PAE-NC (Prenatal Alcohol Exposure-Normal Chow): 10% (v/v) ethanol in non-acidified water *ad libitum* from 10 days prior to mating until gestational days (GD) 8-10. This is intended to replicate typical patterns of drinking during the first trimester of pregnancy in humans. Dams received non-acidified reverse osmosis water for the remainder of pregnancy and normal chow (Rat and Mouse Cubes, Speciality Feeds, Glen Forrest, Australia) throughout pregnancy.
ii. PAE-HMD (Prenatal Alcohol Exposure-High Methyl Donor diet): 10% (v/v) ethanol in non-acidified water *ad libitum* from 10 days prior to mating until GD8-10 and non-acidified reverse osmosis water for remainder of pregnancy. Isocaloric high methyl donor (HMD) chow consisting of 20 mg/kg folate and 4,970 mg/kg choline throughout pregnancy (Speciality Feeds, Glen Forrest, Australia).
iii. H_2_0-NC (Water-Normal Chow): non-acidified water and normal chow throughout pregnancy.
iv. H_2_0-HMD (Water-High Methyl Donor diet): non-acidified water and HMD chow throughout pregnancy.

### Whole-genome bisulfite sequencing of newborn mouse tissues

Pups selected for WGBS in each intervention group were matched on sex and litter size to minimize variability in exposure. Two male and two female pups per treatment group (n = 16 total) were euthanised by intraperitoneal injection with ketamine and xylazine on the day of birth for WGBS of their brain and liver tissues. Mouse tissue samples were stored at -80°C. Remaining littermates grew until adulthood for behavioural testing. Ten milligrams of tissue were collected from each liver and brain and lysed in Chemagic RNA Tissue10 Kit special H96 extraction buffer. Total nucleic acid was extracted from the tissues using the Chemagic 360 instrument (PerkinElmer) and quantified with Qubit DNA High Sensitivity Kit (Catalogue Number: Q32854, Thermo Scientific). 100 ng of genomic DNA was spiked with 0.5 ng of unmethylated lambda DNA (Catalogue Number: D1521, Promega) to assess the bisulfite conversion efficiency. Each sample was digested with 2 µl RNase A (Invitrogen) at 37°C for 20 minutes to remove RNA. 100 ng of genomic DNA from each sample was sheared using a Covaris M220 (300bp settings, Covaris). Libraries were prepared using the Lucigen NxSeq AmpFREE Low DNA Library Kit (Catalogue Number: 14000-1, Lucigen), according to the manufacturer’s instructions. Nextflex bisulfite-seq barcodes (Catalogue Number: Nova-511913, PerkinElmer) were used as the adapters with incubation at 25°C for 30 minutes. The libraries were bisulfite converted using the Zymo EZ DNA Methylation-Gold Kit (Catalogue Number: D5005, Zymo Research) and PCR amplified using the KAPA HiFi Uracil PCR Kit (Catalogue Number: ROC-07959052001, Kapa Biosystems). The final libraries were assessed with the Agilent 2200 Tapestation System using D1000 Kit (Catalogue Number:5067-5582). WGBS was performed by Genomics WA sequencing core on a NovaSeq 6000 (Illumina) using 2×150bp chemistry on an S4 flow cell. The bisulfite conversion rate in each tissue sample was at least 99%. The overall mean coverage in each sample was 9.69x (range: 6.51-12.12x).

### Behavioural testing in adult mice

Littermates who were not sacrificed at birth were reared on normal chow and drinking water *ad libitum* until adulthood (∼8 weeks after birth) when they underwent behavioural tests assessing five neurodevelopmental domains that can be affected by PAE including locomotor activity, anxiety, spatial recognition, memory, motor coordination and balance. These tests included the open field test (locomotor activity, anxiety) (38), object recognition test (locomotor activity, spatial recognition) (39), object in place test (locomotor activity, spatial recognition) (40), elevated plus maze test (locomotor activity, anxiety) (41), and two trials of the rotarod test (motor coordination, balance) (42). Between mouse subjects, behavioural testing equipment was cleaned with 70% ethanol. Video recording was employed for all behavioural tests, except for the rotarod, and the assessment process was semi-automated using ANY-maze software (Stoelting Co., Wood Dale, Illinois, U.S.A.).

### Statistical analysis

Dam characteristics and pup behavioural testing results were generally assessed using unpaired t-tests comparing each treatment group to the baseline control group that was given non-acidified reverse osmosis water and normal chow throughout pregnancy. Trajectories of liquid consumption and weight gain across pregnancy, which were assessed using a quadratic mixed effects model and the trajectory of chow consumption across pregnancy which was assessed using a linear mixed effects model. To examine the effect of alcohol exposure on behavioural outcomes we used linear regression with alcohol group (binary) as the main predictor adjusted for diet and sex. For sequencing data, raw fastq files were mapped to the mm10 mouse reference genome with BSseeker 2 (version 2.1.8) (43) and CG-maptools (version number 0.1.2) (44) using a custom bioinformatics pipeline. CGmap output files were combined as a bsseq object in the R statistical environment (45). We filtered the sex chromosomal reads and then combined reads from mice in the same treatment group using the *collapseBSseq* function, to maximise coverage prior to differential DNA methylation analysis. CpG sites with an aggregated coverage below 10x in each tissue type were removed prior to modelling to ensure there was sufficient coverage in all assessed CpG sites. This retained 94.9% of CpG sites in the brain and 93.8% of CpG sites in the liver. Differentially methylated regions (DMRs) were identified within each tissue using a Bayesian hierarchical model comparing average DNA methylation ratios in each CpG site between PAE and non-PAE mice using the Wald test with smoothing, implemented in the R package *DSS* (46). False-discovery rate control was achieved through shrinkage estimation methods. We declared DMRs as those with a local FDR P-value < 0.05 based on the p-values of each individual CpG site in the DMR, and minimum mean effect size (delta) of 5%. Gene ontology analysis was performed on the brain and liver DMRs using the Gene Set Enrichment Analysis computational method (47) to determine if the DMRs were associated with any transcription start sites or biological processes. Brain and liver DMRs were tested for enrichment within ENCODE Project data sets (48) by an overlap permutation test with 100 permutations using the *regioneR* package. The ENCODE Project data sets that were assessed included ENCFF845WSI, ENCFF764NTQ, ENCFF937JHP, ENCFF269TLO, ENCFF676TSV and ENCFF290MLR. DMRs were then tested for enrichment within specific genic and CpG regions of the mouse genome, compared to a randomly generated set of regions in the mouse genome generated with *resampleRegions* in *regioneR*, with equivalent means and standard deviations. For candidate gene analysis, we compiled a set of key genes and genomic regions identified in previous mammalian PAE studies for site-specific testing based on our prior systematic review (7), which identified 37 candidate genes (Tables S5-6). Murine brain and liver datasets were filtered to candidate gene regions and differential testing was then performed across the entire coding sequence, separately in the brain and liver of the mice on a normal diet using the *callDML* feature in DSS.

### Validation studies in human cohorts

We used existing human data sets to validate observations from our murine model, focusing on regions identified in our early moderate PAE model. Validation studies in human cohorts with existing genome-wide DNA methylation data sets and matching PAE data are described in the Supplementary Material. Briefly, Illumina Human Methylation array .iDAT files were pre-processed using the *minfi* package (49) from the Bioconductor project (http://www.bioconductor.org) in the R statistical environment (http://cran.r-project.org/, version 4.2.2). Sample quality was assessed using control probes on the array. Between-array normalization was performed using the stratified quantile method to correct for Type 1 and Type 2 probe bias. Probes exhibiting a *P*-detection call rate of >0.01 in one or more samples were removed prior to analysis. Probes containing SNPs at the single base extension site, or at the CpG assay site were removed, as were probes measuring non-CpG loci (32,445 probes). Probes reported to have off-target effects in McCartney et al. (50) were also removed. Mouse DMRs were converted into human equivalent regions using an mm10 to hg19 genome conversion with the liftover tool in the UCSC Genome Browser (51). A minimum 0.1 ratio of bases that must remap was specified as recommended for liftover between regions from different species and multiple output regions were allowed. Differential testing of candidate mouse DMRs was carried out using the R package *DMRcate* (52) for each dataset and DMRs were declared as minimum smoothed false-discovery rate (FDR) < 0.05. Cell heterogeneity in each sample including the composition of epithelial, fibroblast and immune cells was estimated from DNA methylation reads using the R package *EpiDISH* (53).

### Data Availability Statement

Data and code pertaining to the original work in this article are available upon request from the corresponding author.

## Author Contributions

D. Martino, M. Symons, A. Larcombe, R. Lister, D. Hutchinson, E. Muggli, J. Craig, J. Halliday, J. Fitzpatrick, S. Buckberry, M. Bestry and E. Elliott contributed to the study design and funding application. E. Chivers, A. Larcombe performed the mouse work including administering the mating, treatments, measurements, monitoring and extracting biological samples. E. Chivers, A. Larcombe and M. Bestry performed the mouse behavioural testing. C. Bakker analysed the videos from the mouse behavioural testing. M. Bestry prepared the whole-genome bisulfite sequencing libraries and performed the data analyses. N. Kresoje assisted with preparing whole-genome bisulfite sequencing libraries. S. Buckberry and D. Martino provided advice and support on the data analysis. J. Halliday and E. Muggli contributed human datasets for reproducibility analysis. M. Bestry and D. Martino drafted the manuscript. M. Symons, A. Larcombe, R. Lister, D. Hutchinson, E. Muggli, J. Craig, J. Halliday, J. Fitzpatrick, S. Buckberry, E. Elliott and N. Kresoje contributed to the development and editing of the manuscript.

## Acknowledgements

We wish to acknowledge the assistance of Dr Jahnvi Pflueger who provided training on preparation of whole-genome bisulfite sequencing libraries. We wish to acknowledge the financial contribution of the Centre for Research Excellence in FASD who supported the murine experiments.

---

**Figure S1.**
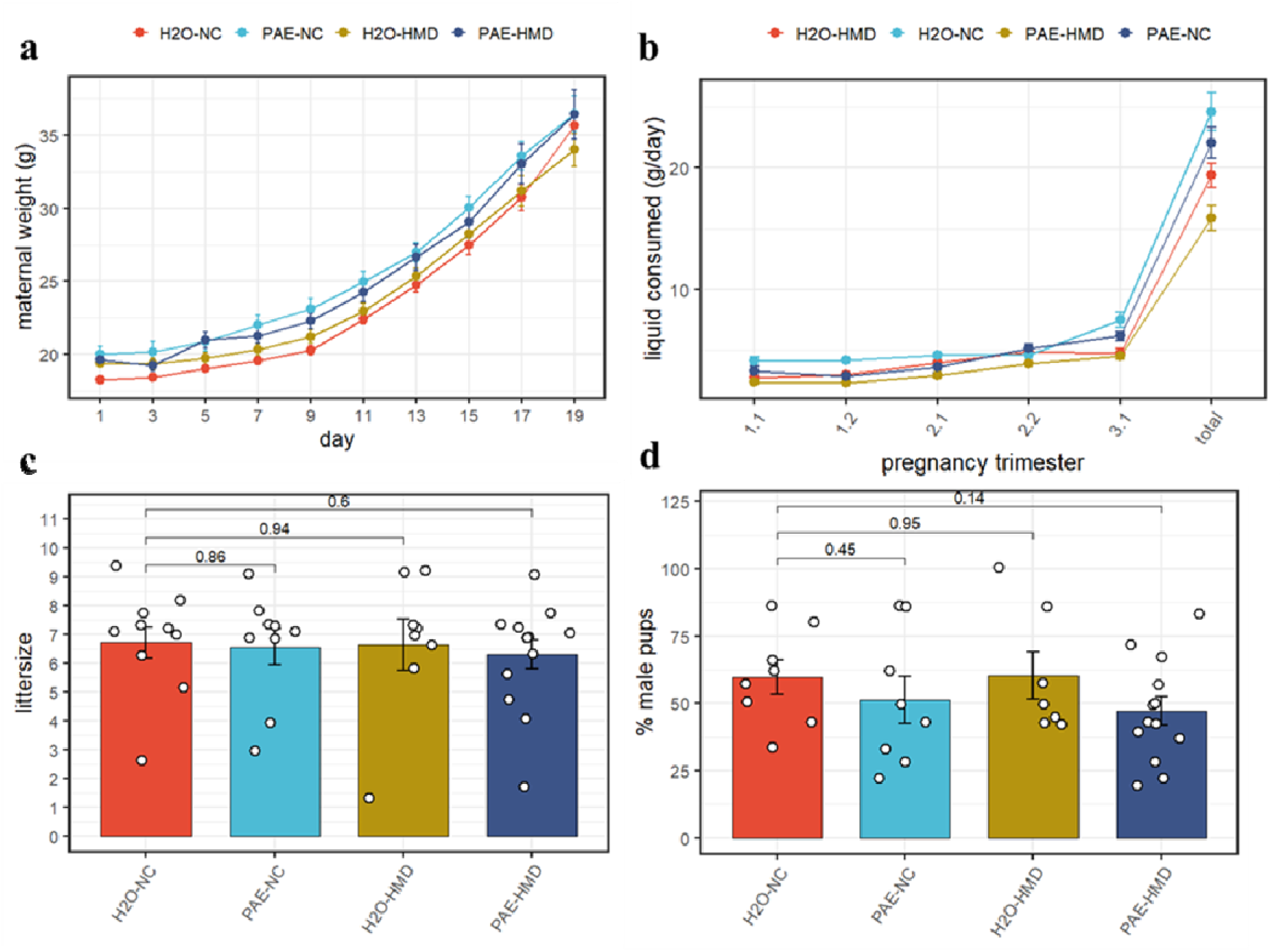
PAE and HMD effects on dam characteristics. (a) Dam weight progression was significantly affected by HMD but not PAE by quadratic mixed-effects model without interaction (b) Trajectory of liquid consumption across pregnancy was affected by PAE and HMD by quadratic mixed effects model. PAE and HMD significantly interacted with trimester of pregnancy. (c) litter size (N = 40) and (d) pup sex ratios (N = 36) were not significantly associated with PAE or HMD by unpaired t-test or ANOVA. All line and bar plots show mean and standard deviation. NC = normal chow, HMD = high methyl diet, PAE = prenatal alcohol exposure. Comparisons show p-value by unpaired t-test compared to the H_2_0-NC baseline treatment group.

**Figure S2.**
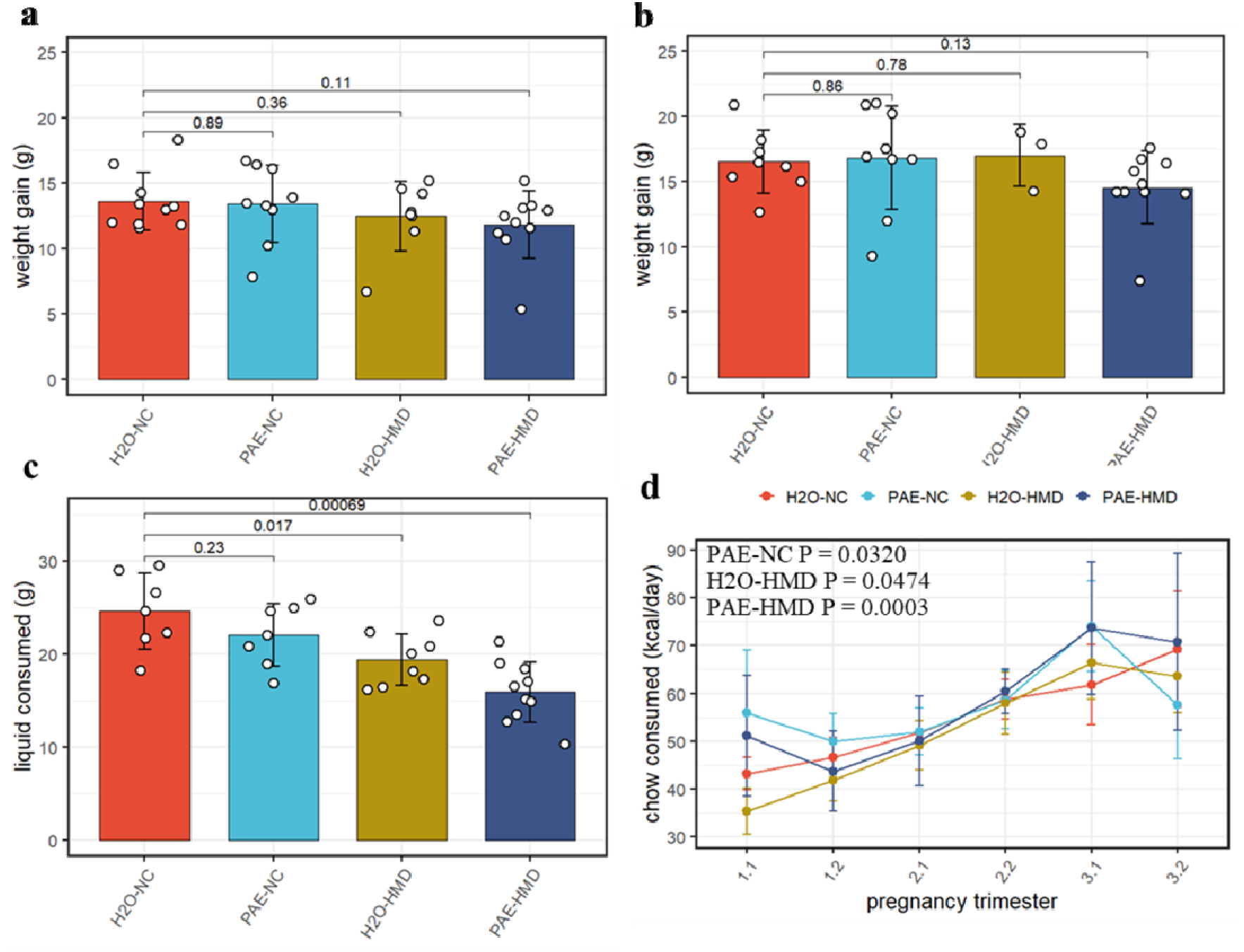
PAE and HMD effects on dam characteristics. There was no significant difference in the average gain of weight in dams between (a) days 1-17 or (b) days 1-19 by treatment group. Both timepoints were included due to some pregnancies ending by day 19. (c) Dams given supplemented chow consumed significantly lower total quantity of liquid across pregnancy. Bar plots show mean and standard deviation for each treatment group. Each point represents one dam. (d) the trajectory of chow consumed by dams across pregnancy significantly varied with the addition of treatments. Points show mean and standard deviation for each treatment group. Statistical analysis involved linear mixed-effects regression comparing trajectories of treatment groups to H2O-NC baseline control group.

**Figure S3.**
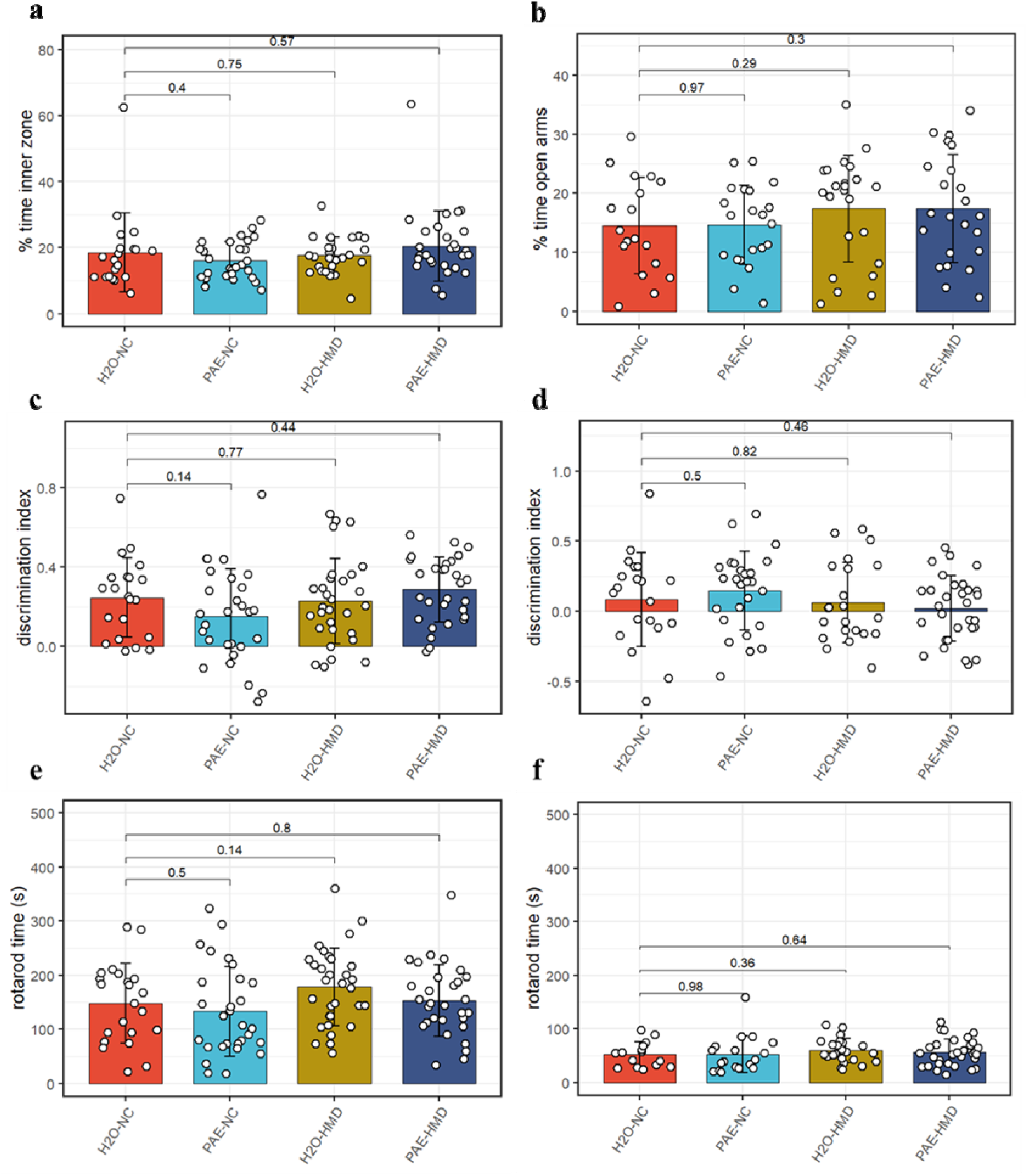
PAE had no significant effect on other assessed behavioural outcomes. PAE and HMD had no significant effect on anxiety as evident by no significant difference by unpaired t-test in the (a) percent time in the inner zone in the open field test (N = 104) and (b) percent time open arms in the elevated plus maze test (N = 85). PAE and HMD had no significant effect on spatial recognition as evident by no significant difference by unpaired t-test in the discrimination index in (c) object recognition (N = 108) and (d) object in place test (N = 98). PAE and HMD had no significant effect on motor co-ordination and balance as evident by no significant difference by unpaired t-test in times in (e) first rotarod test (N = 112) and (f) second rotarod test (N = 87). Bars show mean and standard deviation. Each point represents one mouse. NC = normal chow, HMD = high methyl diet, PAE = prenatal alcohol exposure. Time interval for each mouse was (a-c) 300 seconds and (d) 180 seconds.

**Figure S4.**
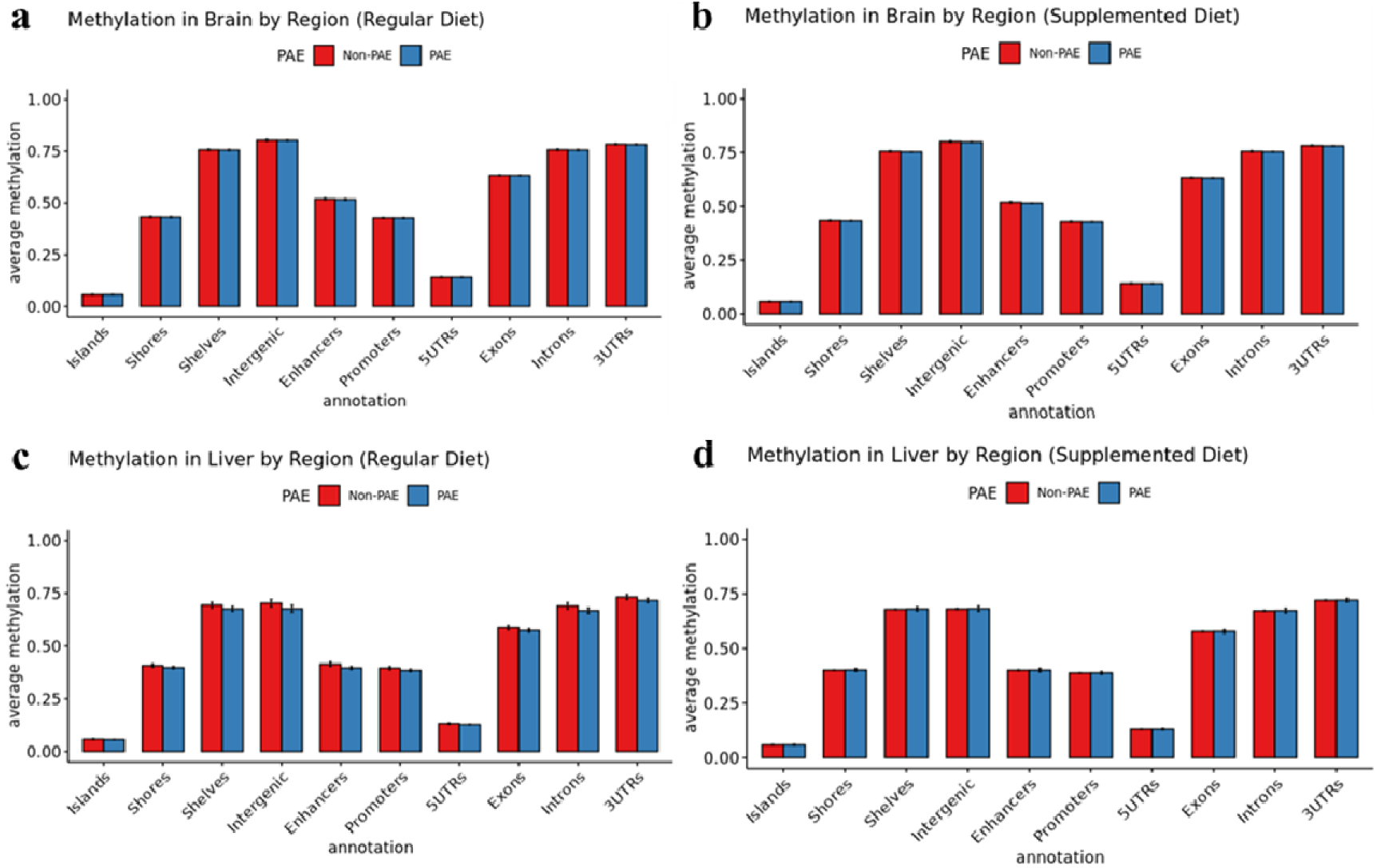
No evidence for global disruption of methylation by PAE. The figure shows methylation levels averaged across CpGs in different regulatory genomic contexts. Neither brain tissue (A & B), nor liver tissue (C & D) were grossly affected by PAE exposure (blue bars). Bars represent means and standard deviation.

